# Cardiorespiratory fitness predicts greater hippocampal volume and rate of episodic associative learning in older adults

**DOI:** 10.1101/578237

**Authors:** Rachel C. Cole, Eliot Hazeltine, Timothy B. Weng, Conner Wharff, Lyndsey E. DuBose, Phillip Schmid, Gardar Sigurdsson, Vincent A. Magnotta, Gary L. Pierce, Michelle W. Voss

## Abstract

Declining episodic memory is common among otherwise healthy older adults, in part due to negative effects of aging on hippocampal circuits. However, there is significant variability between individuals in severity of aging effects on the hippocampus and subsequent memory decline. Importantly, variability may be influenced by modifiable protective physiological factors such as cardiorespiratory fitness (CRF). More research is needed to better understand which aspects of cognition that decline with aging benefit most from CRF. The current study evaluated the relation of CRF with learning rate in the Episodic Associative Learning (EAL) task, a task designed specifically to target hippocampal-dependent relational binding and to evaluate learning with repeated occurrences. Results show that higher CRF was associated with larger hippocampal volume and faster learning rate. Larger hippocampal volume was also associated with faster learning rate, and hippocampal volume partially mediated the relationship between CRF and learning rate. Further, to support the distinction between learning item relations and learning higher-order sequences, which declines with aging but is largely reliant on extra-hippocampal learning systems, we found that EAL learning rate was not related to motor sequence learning on the alternating serial reaction time task. Motor sequence learning was also not correlated with hippocampal volume. Thus, for the first time we show that higher CRF in healthy older adults is related to enhanced rate of relational memory acquisition, in part mediated by benefits on the hippocampus.

## INTRODUCTION

Declining episodic memory is common among cognitively normal older adults, especially after the age of 60 (Craik, 1994; Leal & Yassa, 2015; Nyberg, Lövdén, Riklund, Lindenberger, & Bäckman, 2012). Episodic memory relies on critical hippocampal processes that decline with age (Leal & Yassa, 2015), such as recollection of specific details about an experience, mnemonic discrimination (distinguishing between similar representations), and relational binding (encoding novel relationships between elements of experience) (Konkel & Cohen, 2009; Naveh-Benjamin, 2000). Evidence suggests that declining memory is in part due to negative effects of aging on hippocampal circuits critical for these processes (Driscoll et al., 2003; Geinisman, Detoledo-Morrell, Morrell, & Heller, 1995).

Significant changes in hippocampal structure and function occur in Alzheimer’s Disease (AD) and Mild Cognitive Impairment (MCI) (Jack et al., 2013; Jack et al., 1998; Johnson et al., 2006), yet evidence from both animals and humans demonstrates the hippocampus and associated cognitive functions are also affected during normal aging, even well before observable MCI symptoms (Gallagher & Koh, 2011). For example, older rats have been found to have, among other decrements, worse recollection (Robitsek, Fortin, Koh, Gallagher, & Eichenbaum, 2008), spatial memory (Barnes, 1979), and pattern separation (Burke et al., 2011) than young rats. Similarly, in humans, performance on relational memory tasks that target binding processes (Naveh-Benjamin, 2000; Naveh-Benjamin, Hussain, Guez, & Bar-On, 2003), and mnemonic discrimination tasks that target pattern separation processes (for review see Leal & Yassa, 2018; Reagh et al., 2016) decline with age.

Structurally, tissue loss in the hippocampus occurs during normal aging (Raz, Rodrigue, Head, Kennedy, & Acker, 2004), and this tissue loss can be considered a proxy of lower-level degradation, such as loss of synapses and dendritic complexity. Indeed, declines in hippocampal volume relate to poorer general memory (J. H. Kramer et al., 2007; Mungas et al., 2005), episodic memory (Gorbach et al., 2017; Hedden et al., 2014; Monti et al., 2015), spatial memory (Head & Isom, 2010; Konishi & Bohbot, 2013), relational memory (Etchamendy, Konishi, Pike, Marighetto, & Bohbot, 2012), and mnemonic discrimination (for review see Yassa & Stark, 2011; Yassa et al., 2010). Thus, even otherwise healthy older adults may experience declines in episodic memory processes linked to hippocampal shrinkage.

Notably, there is significant variability in the severity of age-related memory decline experienced between individuals, which may be due in part to differential effects of aging on the physiological and neurobiological processes in the hippocampus (Ash et al., 2016; Gallagher et al., 2006; Rapp & Amaral, 1992; Stark, Yassa, & Stark, 2010; Tomás Pereira, Gallagher, & Rapp, 2015). Critically, individual differences in the trajectory of age-related changes in cognition and neural systems may be influenced by modifiable protective factors such as physical activity (PA) (Hayes et al., 2015; Suwabe et al., 2018; Suwabe et al., 2017) and cardiorespiratory fitness (CRF), which is influenced largely by genetics and PA (Hayes, Forman, & Verfaellie, 2016; Hayes, Hayes, Williams, Liu, & Verfaellie, 2017). CRF is related to brain structure and function (Hayes, Hayes, Cadden, & Verfaellie, 2013) and to better cognitive function broadly (Colcombe & Kramer, 2003; Colcombe et al., 2004; Smith et al., 2010), including episodic memory (Erickson et al., 2009; Hayes et al., 2016; Szabo et al., 2011). In a study that utilized a spatial memory task, Erickson et al. (2009) found that hippocampal volume mediated the relationship between CRF and memory in older adults. Results from exercise training interventions have further shown that change in CRF relates to change in hippocampal volume (Erickson et al., 2011) and cerebral blood flow (Maass et al., 2015).

However, these studies supporting a relationship between CRF, hippocampal structure, and memory in older adults have primarily used spatial working memory and spatial object recall and recognition tasks. Although these tasks target some aspects of hippocampal function (e.g., spatial memory), they also emphasize one-trial learning, and while one role of the hippocampus is to acquire relations from single episodes at a time (Henke, Buck, Weber, & Wieser, 1997), the hippocampus is also involved in actively maintaining novel information over short time periods (Ranganath & D’Esposito, 2001; Watson, Voss, Warren, Tranel, & Cohen, 2013) and dynamically integrating information that connects episodes over time (Koster et al., 2018). One-trial learning does not capture this accumulation of relations over repeated occurrences with overlapping content, which requires discriminating between similar memories (e.g., seeing Bill at two coffee shops) while also accessing and strengthening the relationships between these experiences (e.g., Bill) with repeated occurrences. Because aging is known to impair relational binding (Naveh-Benjamin, 2000) and discrimination of similar memories (Leal & Yassa, 2015), a task tapping into the ability to rapidly build distinct, but similar, relational memories should be maximally sensitive to hippocampal circuits affected early in aging but spared with higher CRF.

In this vein, we designed the Episodic Associative Learning (EAL) task to examine paired associates learning (Figure 1). The task measures the rate of learning item pairs with overlapping elements (e.g., A-B, A-C), for which we reliably observe strong age differences (Clark, Hazeltine, Freedberg, & Voss, 2018). Overlapping elements require each pair to be kept distinct as participants learn via trial and error. A rapid learning rate represents the ability to quickly form similar but distinct relations, which should theoretically reflect hippocampal processes of mnemonic discrimination and relational binding. However, we have not previously tested our prediction that faster learning rate is related to hippocampal integrity in older adults, which is the first goal of this study. We also evaluate our prediction with the striatum (caudate and putamen) as control regions, as these are subcortical regions that also deteriorate with aging (Raz et al., 2005a; Raz et al., 2003) and are involved in extracting regularities of experience across time (Poldrack & Packard, 2003; Seger, 2006). Second, we test the prediction that higher CRF is related to learning rate and the extent to which this is mediated by hippocampal (but not striatal) volume. Further, to distinguish between outcomes of learning item relations and higher-order sequences over time, we compare the relation of CRF with EAL rate to motor sequence learning in an alternating serial reaction time task (ASRT). ASRT performance has been shown to decrease with age (Howard & Howard, 1997), but also to depend more on the striatum than the hippocampus (for review see Howard & Howard, 2013).

**Figure 1.**
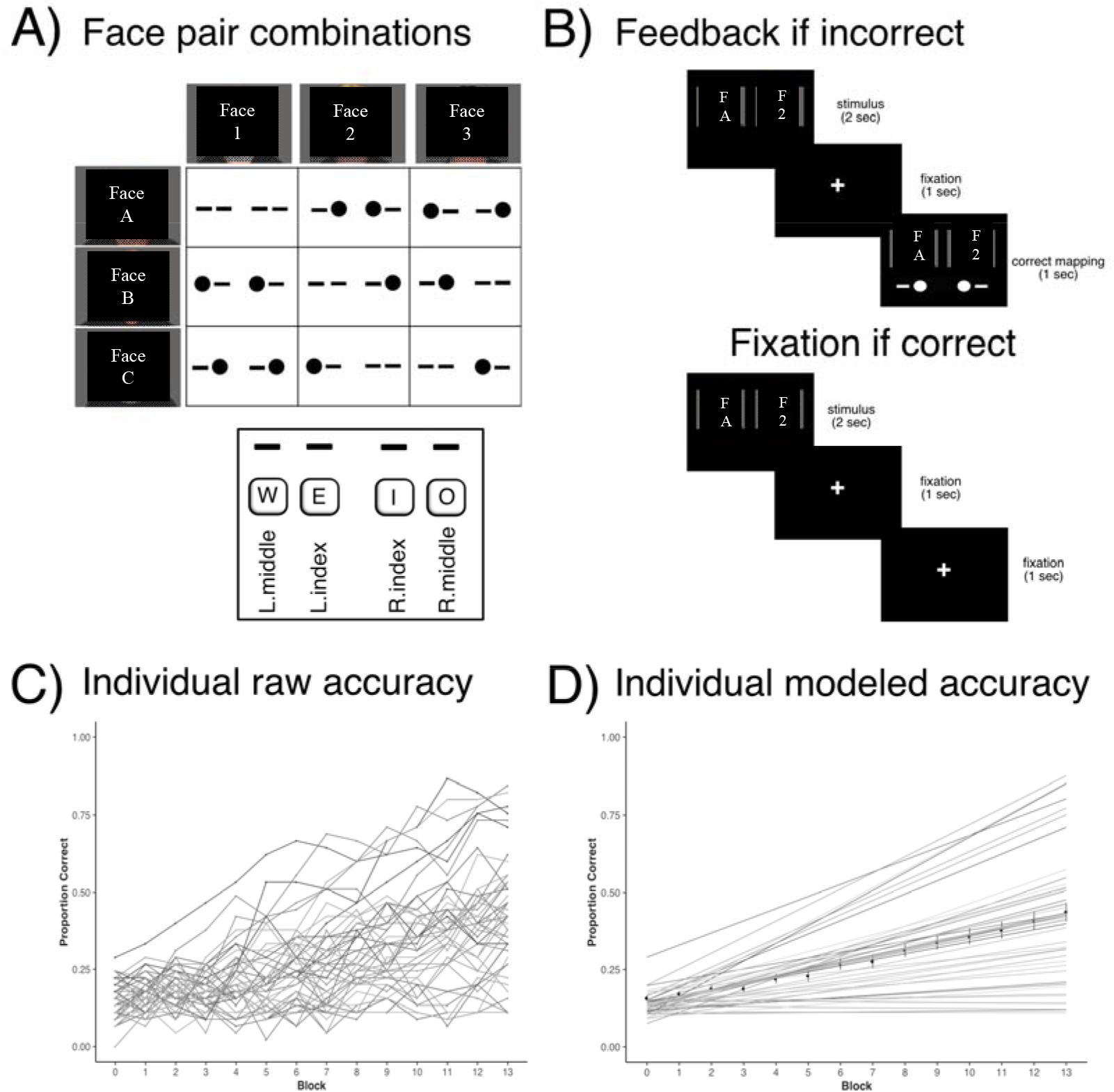
A) Layout of face-face objects pairs and the corresponding keypress(es). Numbers and letters represent individual male faces. B) Incorrect and correct example trials. In the feedback, filled-in circles represent a keypress in the key that corresponds to the location of the circle, and dashes represent no keypress for that key. C) Raw data for all participants from EAL task. D) Data modeled by linear model for all participants, with average represented by thick black line.

Still, other cognitive processes that decline with aging, such as processing speed and working memory (Salthouse, 1994), may play a role in EAL performance. It is possible that slow processing and poor working memory could contribute to a slow learning rate on the EAL task. However, in our previous study (Clark et al., 2018), we found that while processing speed was related to EAL performance, it did not account for the age differences. Here we aim to replicate the effect for processing speed, and we further evaluate whether working memory accounts for relationships between CRF and the EAL learning rate.

Results from this study will provide a more comprehensive understanding of how age-sensitive learning processes correspond to individual differences in brain structure and modifiable lifestyle health characteristics.

## METHODS and ANALYSES

### Participants

Participants were older adults recruited from Iowa City and the surrounding communities (see Table 1). Participants were recruited using approved University email advertisements, local fliers and approved advertisements at the University of Iowa Hospitals and Clinics. Eligibility for all participants required the following criteria: 1) have no self-reported psychiatric and/or neurological condition, including depression, anxiety disorder, ADD or ADHD, epilepsy, meningitis, Parkinson’s disease, stroke, brain surgery, and head injury; 2) have no diagnoses of any of the following conditions: heart condition or other cardiovascular event, COPD, uncontrolled asthma (not on medication or inhaler for the past three months or more), cystic fibrosis, unregulated thyroid disorder (not on medication for the past 3 months or more), renal or liver disease, heart murmur, and smoking or living with someone who smokes in the past 3 months; 3) have normal color vision; 4) have corrected visual acuity of 20/40 or above; and 5) have no self-reported regular use of steroid-based medication, psychotropics, recent or current chemotherapy, or medications that indicated diagnosis of a chronic psychiatric disorder. All participants provided written informed consent approved by the University of Iowa Institutional Review Board (IRB). All study procedures were in accordance with the University of Iowa IRB’s policies. All participants were screened using the Mini-Mental State Examination (MMSE) and were excluded if they scored less than 24 points (out of 30).

The full sample consists of 45 participants aged 60-80. Thirty-seven participants were low-active, self-reporting < 30 minutes of moderate intensity activity twice a week. Eight participants were highly-active, self-reporting performing moderate to vigorous PA for 5 or more days per week (on average) for 45 minutes per session for at least the past 2 years or longer. All participants completed at least 5 laboratory visits. Participants first attended an orientation session that included reviewing the IRB form, obtaining consent, and completing cognitive screening, health history, and self-report questionnaires, as well as a mock MRI in a simulator. The second visit included the maximal exercise test. The third and fourth visits consisted of extensive cognitive testing, with each visit lasting about 2 hours. The final visit was the MRI session, which included structural and functional scans. A subset of participants was from the pre-intervention sessions of an exercise intervention (NCT02453178).

**Table 1.** Sample demographics

|  | **N** | **Age**<br>**Range** | **Age**<br>**Mean (SD)** | **Years Education**<br>**Mean (SD)** | **MMSE**<br>**Mean (SD)** |
| --- | --- | --- | --- | --- | --- |
| *Total* | 45 | 60 - 76 | 66.51 (4.24) | 18.13 (2.8) | 29.18 (1.17) |
| Male | 16 | 60 - 76 | 67.12 (5.04) | 18.56 (3.39) | 28.88 (1.20) |
| Female | 29 | 60 - 74 | 66.17 (3.77) | 17.90 (2.45) | 29.34 (1.14) |

### Cardiorespiratory Fitness Testing

Maximal oxygen uptake was measured with indirect calorimetry using a maximal exercise test on a cycle ergometer with resistance increasing in two-minute intervals. Oxygen consumption was calculated from expired air samples at 15-s intervals until peak VO_2_ was reached. VO_2_max was determined and test terminated when a) respiratory exchange rate (RER) exceeded 1.10, b) participant reached 90% of age-predicted heart rate maximum, or c) heart rate and/or oxygen uptake plateaued despite an increase in resistance level. This test was also terminated if the participant showed signs of distress or if physiological signals became abnormal (blood pressure, heart rate, EKG). VO_2_max is the gold standard for measuring CRF. Because participants were over the age of 40, a physician was present during the testing. In all analyses we use relative VO_2_max (mL/kg/min) to adjust for weight.

### Cognitive testing

Multiple tasks were administered to measure relational learning and memory (EAL), motor sequence learning (ASRT), working memory (Face N-back), and processing speed (Pattern and Letter Comparison).

#### Episodic Associative Learning

This task has previously been described fully in Clark et al. (2018) (Experiment 2). Briefly, participants learned to respond to face pairs with either bimanual, unimanual, or no key press (Figure 1A). The correct keypress was unique for each face pair. Importantly, no single face provided information about the correct keypress.

To familiarize participants with the task, a brief practice phase (five minutes) occurred immediately before the learning phase with practice stimuli (animals and modes of transportation) and a slower pace of trials. In each trial, participants were shown a stimulus pair in the center of the screen and were to respond with their middle and index fingers on keys W, E, I, O, respectively. For each trial, if the participant responded correctly, the next trial began following a fixation screen with a centered fixation cross. If the participant responded incorrectly, however, the pair was presented again with the correct mapping shown directly below the stimuli (Figure 1B). Practice was followed by a learning phase including 14 blocks of face pairs. Each of the nine pairs appeared five times in each block in a randomized order. Face stimuli were chosen from young and older adult male neutral faces in the Center for Vital Longevity Face Database. For each participant, either young or older faces appeared on the left and the other age category of faces appeared on the right. Category (young and old faces) and position (left and right) was counterbalanced between participants. Between each block, participants received feedback regarding accuracy and speed of response of each hand. The next block began when the participant decided to continue. The task lasted for approximately 50 minutes.

#### EAL Analysis

Raw accuracy data was fit to a linear mixed effects model using R (Figure 1C and 1D). Mixed effects modeling was selected over repeated-measures ANOVA to more accurately model individual differences in learning rate. Specifically, we modeled a linear slope parameter to index learning rate across blocks and a quadratic parameter to index change in learning rate. The model was fit using R’s linear mixed-effects (lme4) package (Pinheiro & Bates, 2000), which simultaneously estimates all fixed and random effects using maximum likelihood estimate. We began with the most complex model for each analysis and compared simpler models using model-comparison procedures based on the Bayesian Information Criterion (Schwarz, 1978). The final model and parameter estimates for the best-fitting model are described in the results. The primary variable of interest in our analyses was learning rate based on the linear slope parameter for each individual, which represents the speed at which the individual acquired the correct mappings for the face pairs.

#### ASRT

In the ASRT, four open circles were in the center of the computer screen. On each trial, one of the four circles became filled in, and the participant was instructed to press the corresponding key (using their left and right middle and index fingers) as quickly and accurately as possible. The circle remained filled until the participant pressed the correct key, which immediately initiated the next trial in which a different circle was filled in. On alternating trials, the circles followed a specific sequence (sequences were counterbalanced across participants), while on the other trials, they were randomly ordered, with the constraint that no trial could repeat the circle from the previous trial. Participants completed 32 blocks with 90 trials each, for a total of 3,072 trials. Blocks were separated by a mandatory 30-second break. A subset of participants (36 of 45) completed this task.

#### ASRT Analysis

Inaccurate trials were discarded from analysis. The difference score for each block was calculated as the difference between average RT to sequence trials and random trials, with higher difference scores representing faster responses to sequence items. For the main dependent variable, we averaged the difference scores for the blocks in the last quarter of the task (blocks 25-32), where performance seemed to plateau.

#### Face N-back

The face N-back task was administered in the scanner and consisted of participants viewing a continuous stream of neutral faces. For each face, participants were asked to determine whether each face matches the face presented *n* items before. This task included 1-back and 2-back conditions, with the 2-back being most demanding on working memory. Data was collected for the Face N-back task for 42 of the 45 participants. The data for one subject experienced a technical issue, and 2 subjects had discomfort because of large head size, thus the N-back task was skipped to accommodate modified scan time.

#### Face N-back Analysis

Reaction time and accuracy are the primary performance outcomes. The average difference in accuracy between the 1-back and 2-back blocks represents a working memory cost. Since participants were encouraged to respond accurately and reaction time was less emphasized, we use accuracy cost as the main dependent variable.

#### Processing Speed

Participants completed a paper-and-pencil pattern and letter comparison task (Salthouse, 1996). The pattern comparison section contained two trials, each consisting of 30 pairs of line drawings (“patterns”). The participant compared the patterns and wrote either “S” (same) or “D” (different) on a line between the items for as many items as possible within 30 seconds. The letter comparison section consisted of two trials, each consisting of one page with 15 pairs of letter strings. Again, the participant was to write S or D on a line between the letter strings for as many items as possible within 30 seconds.

#### Processing Speed Analysis

We calculated the average number of correct pattern or letter string pairs completed between the two trials of each, normalized across participants within the pattern and letter sections, and then averaged the two z-score values to get a processing speed score for each person.

### Magnetic Resonance Imaging

#### MRI Protocol

All magnetic resonance imaging (MRI) was conducted at the Magnetic Resonance Research Facility (MRRF) at University of Iowa. Toward the beginning of data collection, in June 2016, MRRF retired a Siemens scanner and acquired a new GE scanner. Thus, MRI data were acquired with either a 3.0T MRI Siemens TIM Trio scanner using a 12-channel head coil (N = 20), or 3.0T General Electric (GE) Discovery MR750w MRI Scanner using a 32-channel head coil (N = 25). For the scans collected on the Siemens scanner, a three-dimensional magnetization-prepared rapid gradient echo (MPRAGE) T1 scan was collected with the following parameters: echo time (TE)=3.09ms, repetition time (TR)=2530ms, inversion time (TI)=900ms, flip angle=10°, Acquisition Matrix=256 x 256×240mm, Bandwidth=219 Hz/pixel, voxel dimensions=1.00 x 1.00 x 1.00, number of slices=240. For the scans collected on the GE scanner, a three-dimensional fast spoiled gradient echo sequence (FSPGR) T1 scan was collected with the following parameters: TI=450ms, TE=3.376, TR=8.588ms, flip angle=12°, Acquisition Matrix=256×256×240, FOV=256×256×240, voxel dimensions=1.00 × 1.00 × 1.00, number of slices=240.

#### Analysis

Subcortical volume estimates were calculated using Freesurfer’s automated subcortical segmentation tool (Fischl et al., 2002; Fischl et al., 2004). Subcortical volumes were adjusted based on intracranial volume (ICV) to account for individual differences in body size and gender (Raz et al., 2005b). Adjustment was performed separately for each region for each hemisphere using the formula based on the analysis of covariance approach: adjusted volume = raw volume – *b*(ICV – mean ICV), where *b* is the coefficient of regression of the region volume on ICV, and ICV is the participant’s total ICV estimated via Freesurfer. For each region, the adjusted left and right hemisphere values were summed to provide an average bilateral volume. Separate hemisphere volumes and bilateral volumes for regions of interest (hippocampus, caudate, putamen) were used in the analyses.

## RESULTS

See Table 1 for participant demographics and Table 2 for descriptive statistics for variables of interest.

**Table 2.** Descriptive statistics for variables of interest

|  | **CRF**<br>**Mean (SD)** | **Hippocampal**<br>**volume**<br>**Mean (SD)** | **Caudate**<br>**volume**<br>**Mean (SD)** | **Putamen**<br>**Volume**<br>**Mean (SD)** | **EAL Learning**<br>**Rate**<br>**Mean (SD)** |
| --- | --- | --- | --- | --- | --- |
| *Total* | 24.01 mL/kg/min<br>(7.69) | 6365.69 mm<sup>3</sup><br>(666.17) | 7066.33 mm<sup>3</sup><br>(721.28) | 9297.70 mm<sup>3</sup><br>(889.64) | .02 proportion<br>per block (.02) |
| Male | 26.73 (7.76) | 6607.55<br>(665.00) | 6976.69<br>(683.51) | 9425.04 (853.90) | .02 (.02) |
| Female | 21.29 (7.06) | 6123.82<br>(677.65) | 7152.98<br>(745.53) | 9170.35 (910.62) | .02 (.01) |

### Hippocampal volume is associated with faster learning on EAL task

We first tested whether hippocampal volume predicts how quickly individuals learned the face pairs and their associated responses. Consistent with predictions, individuals with larger hippocampal volume more quickly acquired the correct responses (r = .36, p = .02) (Figure 2). This was observed for both hemispheres (right: r = .35, p = .02; left: r = .35, p = .02). Neither age, sex, nor years of education were related to hippocampal volume or EAL slope, so no covariates were included in this analysis.

**Figure 2.**
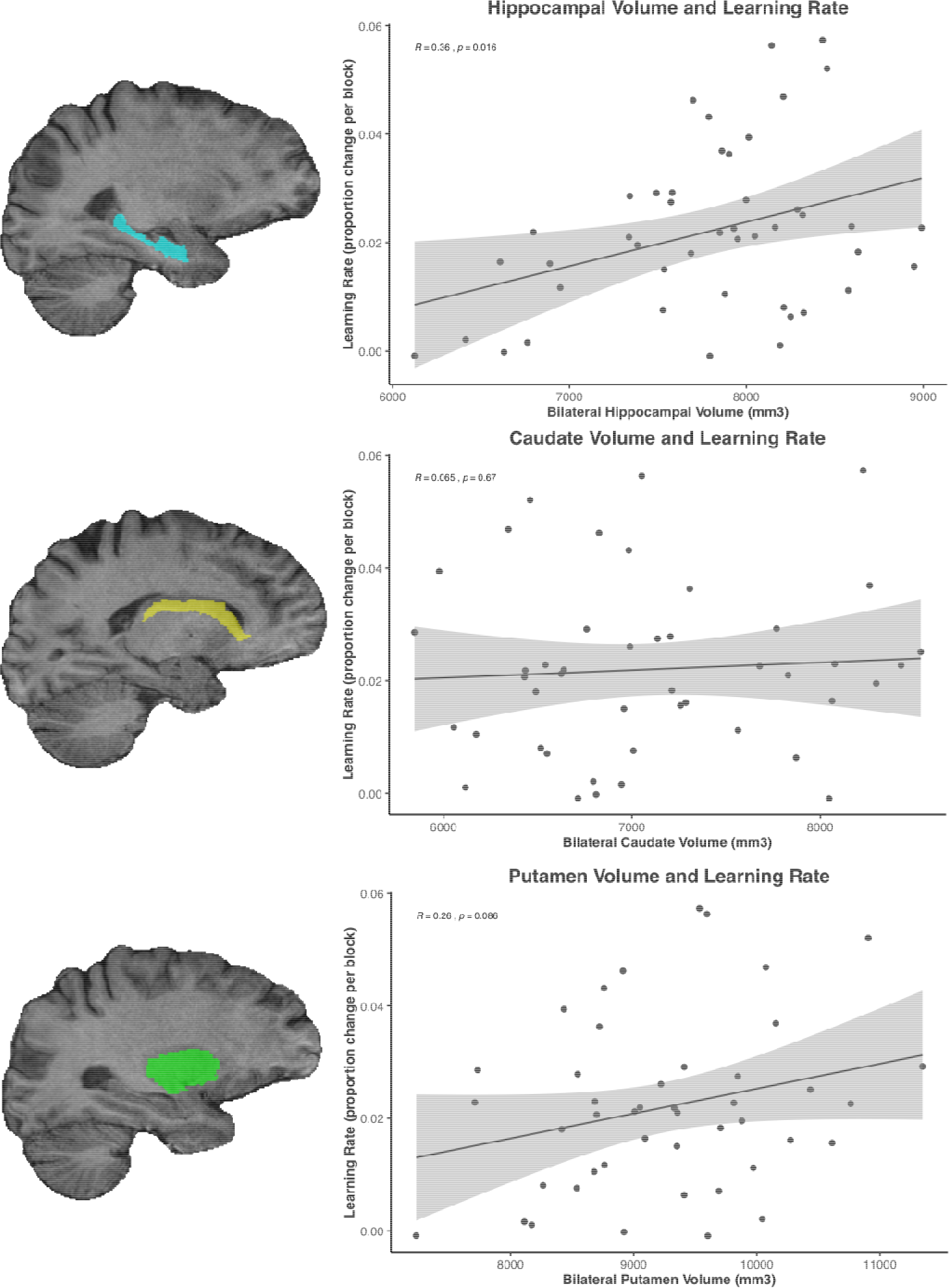
Example subject-specific masks for right hippocampus, right caudate, right putamen from coronal slices. Scatterplots showing relationships between bilateral hippocampal volume, bilateral caudate volume, bilateral putamen volume, and EAL learning rate.

Unlike hippocampal volume, volume of neither the cauduate (r = .06, p = .67) nor putamen (r = .26, p = .09) correlated with learning rate (see Figure 2). For both structures, the relationship of bilateral volume with learning rate did not differ from the relationship of left and right hemisphere volume with learning rate. These findings support that EAL learning rate is uniquely related to hippocampal volume rather than being related to other subcortical regions involved in motor learning. The results suggest EAL learning rate involves processes that extend beyond statistical regularities over time.

To evaluate the relationship between different types of learning, we also examined learning rate on the ASRT. Participants demonstrated sequence learning as, on average, responses became faster over time to the sequenced relative to random items (based on average difference score for last 8 blocks of task, t(36) = 3.1, p = .002). To support their distinction, EAL rate was not correlated with ASRT learning (r = .11, p = .51). Further, ASRT learning was not correlated with hippocampal volume (bilteral: r = .002, p = .1; right: r = .02, p = .9; left: r = −.01, p = .9). Surprisingly, though, ASRT learning was also not correlated with caudate (bilateral: r = −.28, p = .10; right: r = −.26, p = .12; left: r = −.27, p = .11), or putamen (bilateral: r = −.09, p = .60; right: r = −.14, p = .42; left: −.03, p = .84) volume. These results provide additional evidence that EAL performance is uniquely tied to hippocampal processes.

To control for possible effects of processing speed and working memory, we tested whether the average processing speed measure from the Pattern and Letter Comparison tasks and the cost measure from the Face N-back task were related to EAL rate and hippocampal volume. We found that EAL rate was not related to average processing speed (r = .23, p = .13), but was related to working memory cost (r = −.34, p = .03), such that individuals who learned faster on the EAL also had less cost on the face N-back task. However, bilateral hippocampal volume was not correlated with processing speed (r = .08, p = .6), or with working memory cost (r = .11, p = .51). Further, the relationship between hippocampal volume and EAL rate remains significant when including processing speed (t = 2.4, p = .02) and working memory cost (t = 2.5, p = .02) as covariates. Thus, neither processing speed nor working memory accounts for the relationship between hippocampal volume and EAL learning rate.

### Cardiorespiratory fitness positively predicts learning rate on EAL task

Next, we tested whether CRF predicted EAL learning rate, and whether hippocampal volume mediated this relationship. Although CRF was related to sex (point-biserial correlation: r = −.35, p = .02), since learning rate was not related to sex (r = −.03, p = .85), we did not include sex as a covariate. Similarly, although age was related to CRF (r = −.37, p = .01), age was not related to learning rate (r = −.20, p = .20), so age was not included as a covariate. Lastly, years of education was related to CRF (r = .34, p = .02), but not to learning rate (r = .16, p = .29). Thus, no covariates were used in this analysis. As predicted, individuals with higher CRF had a faster EAL learning rate (r = .45, p = .002). CRF also had a positive but non-statistically significant relationship with processing speed (r = .28, p = .06). However, since processing speed was not related to learning rate on the EAL task, processing speed could not account for the relationship between CRF and EAL learning rate. Additionally, CRF was not related to working memory cost (r = −.03, p = .88), which also discounts the possibility that working memory is a confound in the association between CRF and EAL learning rate.

### Relation of cardiorespiratory fitness with hippocampal volume

CRF was related to hippocampal volume with a positive trend (r = .25, p = .09, Figure 3). Though the relationship was not statistically significant, the effect size of r = .25 is similar to what has been found in different samples of older adults (Erickson et al., 2009). This relationship was slightly stronger for the left hippocampus than the right (left: r = .28, p = .07; right: r = .21, p = .15). As comparison, the correlations of CRF with bilateral caudate volume (r = .03, p = .8) and bilateral putamen volume (r = .01, p = .96) were both non-significant and near zero effect-size.

**Figure 3.**
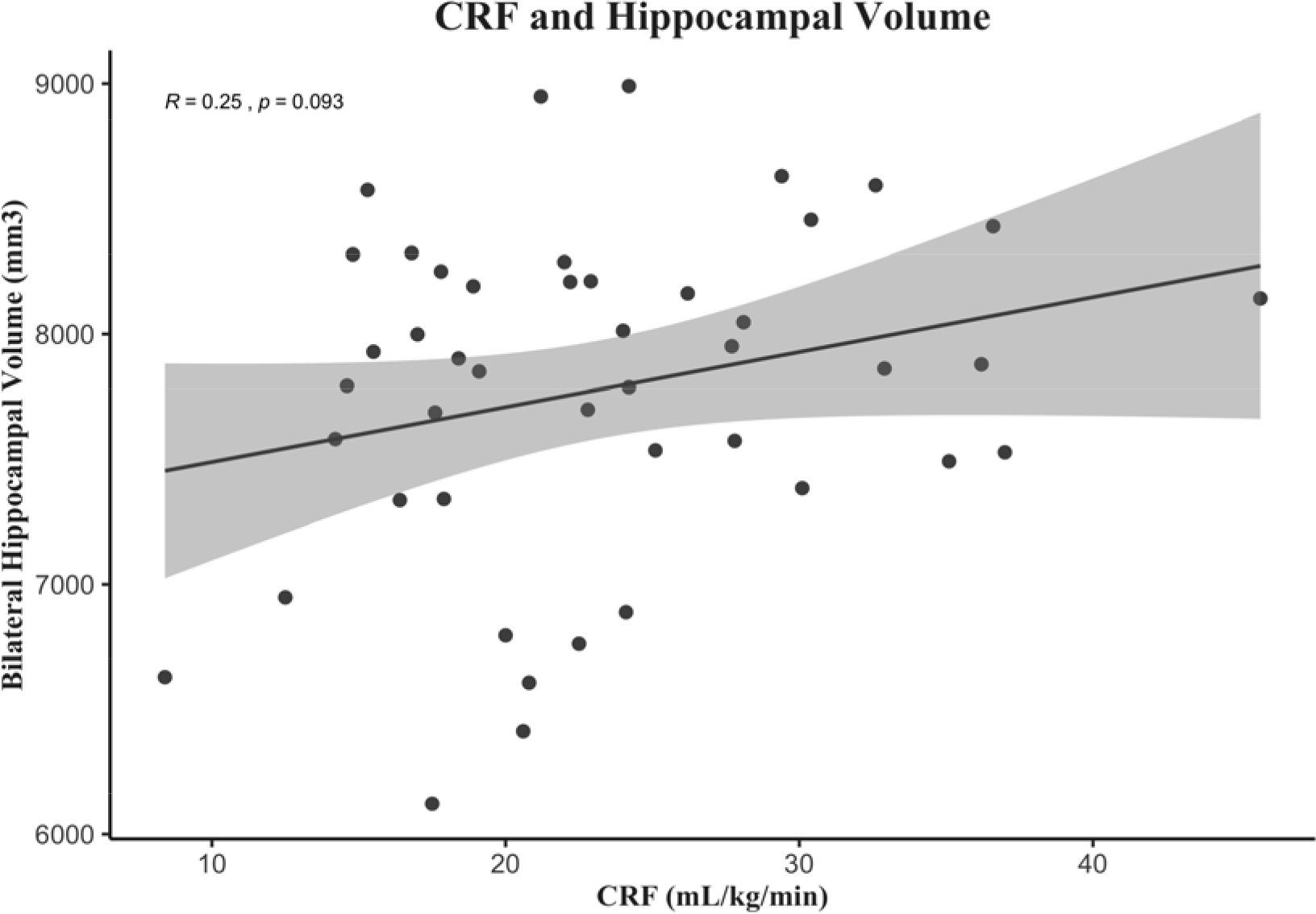
Relationship between CRF and bilateral hippocampal volume.

### Hippocampal volume partially mediates the relationship between CRF and learning rate on EAL task

Because CRF positively predicted both hippocampal volume and learning rate, and hippocampal volume was positively associated with EAL learning rate, we ran a mediation model to test whether hippocampal volume acts as a mediator in this relationship. Based on the lack of contribution of age, sex, and education to the relationships between variables of interest, no covariates were included in the mediation model. The steps outlined by Baron and Kenny (1986) were completed in R using the *mediate* package.

Results indicate that hippocampal volume partially mediates the relationship between CRF and learning rate. The first step of setting up the mediation is to test for the direct effect of the independent variable on the dependent variable. As previously shown, the total effect of CRF on learning rate was significant (ß(43) = .45 (SE = .14), t = 3.3, p = .002). The direct effect of CRF on learning rate after taking hippocampal volume into account as a mediator was reduced but was still significant (ß(42) = .39, (SE = .14), t = 2.8, p =.007). The final and critical step of mediation is to determine whether the indirect effect (i.e., the amount that hippocampal volume mediates the relationship) is significant. We tested the indirect effect using a bootstrap estimation approach with 5000 simulations, which revealed that the indirect effect was nearly, but not statistically, significant (b = .07, 95% CI [−.002, .18], p = .06).

## DISCUSSION

In the current study, we found CRF positively predicted learning rate on the EAL task, which we designed to heavily draw on hippocampal processes that decline with age. We found a specific relationship of EAL task performance with hippocampal volume, such that rate of learning was related to hippocampal volume but not volume of other sub-cortical structures involved in learning (e.g., caudate and putamen). Critically, hippocampal volume was related to both CRF and learning rate on the EAL task and served as a partial mediator of this relationship. These findings provide evidence that CRF, which is a modifiable physiological characteristic associated with hippocampal structure and function, serves as a modifier of age-related changes in hippocampal structure and associated episodic memory acquisition.

Our finding of a relationship between CRF and episodic learning rate aligns with previous findings of positive relationships between CRF and cognitive outcomes (Erickson et al., 2009; Hayes et al., 2016; Hayes et al., 2013; Hayes et al., 2017; Szabo et al., 2011), and is supported by a large body of work in animals and humans that has shown beneficial effects of PA on the brain, specifically the hippocampus, and hippocampal-dependent learning and pattern separation (Creer, Romberg, Saksida, van Praag, & Bussey, 2010; Hayes et al., 2015; Suwabe et al., 2017; van Praag, Shubert, Zhao, & Gage, 2005). Critically though, previous work linking CRF to hippocampal volume and cognition in older adults has not examined performance on tasks that require acquisition of object-based relations over time, a process that may be more sensitive to subtle aging processes in the hippocampus (Reagh et al., 2016). The EAL task uniquely addresses this gap by providing a measure of acquisition rate across repeated co-occurrences of paired elements. We have previously shown robust age differences in performance on this task (Clark et al., 2018), which suggests that it is sensitive to processes that change during cognitively normal aging. The task draws on relational binding and mnemonic discrimination for the acquisition of overlapping but distinct relations over repeated co-occurrences. To target mnemonic discrimination processes, the relations used in the EAL task have overlapping elements across pairs but also require distinct representations in order to learn the unique responses. For these reasons, the EAL task is a useful tool for examining subtle age-related differences, and the current study revealed a novel relationship between CRF and learning rate. This study also provides validation that the EAL task is a useful tool for targeting associative bindings (see Supplementary materials), and the addition of multiple data points throughout the task allows researchers to model acquisition curves and more thoroughly probe binding processes over repeated occurrences of item pairs.

We also found that learning on the EAL task was distinct from other aspects of cognition. EAL task learning rate was not related to sequence learning, which has been shown to be less affected by age than the learning processes involved in the EAL task. Sequence learning was also not related to hippocampal volume. We further tested whether working memory and processing speed could account for the relationships between CRF, hippocampal volume, and learning rate. As working memory and processing speed are known to decline with age, it is possible that these age-related changes could influence performance on the EAL task. In our previous study (Clark et al., 2018), we found that individual differences in processing speed did not account for differences in EAL learning rate. We did not, however, measure working memory in the previous study. In the current study, we again found that processing speed was not related to performance on the task, nor was it related to hippocampal volume or CRF. We also found that although working memory was related to EAL learning rate, it was not related to hippocampal volume or CRF. Thus, the results suggest working memory cannot account for the relationships between CRF, hippocampal volume, and EAL learning rate. This finding of specificity is consistent with, and extends upon, our previous findings. Combined, our previous findings and the current findings provide evidence for a specific relationship between CRF, the hippocampus, and the learning processes involved in the EAL task.

The EAL paradigm involves learning unique responses to overlapping pairs of stimuli, thus relying on hippocampal circuits to create and continuously strengthen distinct bindings. The multiple, interleaved presentations of each pair require rapid binding, and continuous maintenance, updating, and integration of relations, processes supported by the hippocampus (Henke et al., 1997; Ranganath & D’Esposito, 2001). Koster and colleagues (2018) recently provided evidence of a big-loop recurrence process within the hippocampus that allows it to both store representations of distinct episodes and integrate information across related episodes, which together would support richly constructive yet precise episodic memories. Koster et al. (2018) found that big-loop recurrence could account for successful inference, which consists of binding information that does not exist simultaneously but does have common relations across episodes. While our task does not rely on inference in this same sense, rapid learning requires managing overlapping yet distinct relations across trials. Future research could use the EAL task and ultra-high-resolution imaging to test whether the activity of hippocampal circuits during rapid learning is similar to the recirculation of hippocampal inputs that supports inference.

Hippocampal volume was used in the current study as a biomarker of hippocampal integrity. As such, measuring hippocampal volume has many strengths. Automated segmentation of the hippocampus is quite robust (Fischl et al., 2002), has been widely used, and does not require manual corrections or decisions. Hippocampal volume has also been shown to be a sensitive biomarker to other health factors (Erickson et al., 2011; Kleemeyer et al., 2016; for review see Ott, Johnson, Macoveanu, & Miskowiak, 2019). Nevertheless, volumetric measurements do involve some limitations. As the hippocampus is far from a homogenous region, structurally or functionally, examination of bilateral hippocampal volume as measured from the entire hippocampus may be too broad. Findings within the last few decades have supported the differential roles of distinct regions within the human hippocampus (for review see Poppenk, Evensmoen, Moscovitch, & Nadel, 2013), specifically specialization along the longitudinal-axis. The hippocampus can be generally divided into an anterior (ventral in rodents) and posterior (dorsal in rodents) portion. Of relevance for the current study, in humans, the anterior hippocampus has been found to be involved in encoding of novel stimuli, whereas the posterior region is known for its involvement in spatial processing (Poppenk et al., 2013; Ryan, Lin, Ketcham, & Nadel, 2010; Woollett & Maguire, 2011). Regarding PA and CRF in relation to anterior and posterior sections of the hippocampus, the results are mixed, but generally favor the anterior hippocampus. Low-intensity walking has been found to be more strongly correlated with anterior hippocampus compared to posterior hippocampus (Varma, Tang, & Carlson, 2016), and a 1-year aerobic exercise intervention selectively increased the volume of the anterior, but not the posterior, hippocampus (Erickson et al., 2011). In the same study by Erickson et al. (2011), change in CRF was related to increases in both anterior and posterior hippocampus. Both Maass et al. (2014) and Thomas et al. (2016) found a specific relation between increase in CRF and increase in anterior hippocampal volume, suggesting greater sensitivity to CRF in the anterior hippocampus. However, sub-regional measurements do have methodological challenges. As automation is not as robust for sub-regions in 3T, decisions must be made as to how to segment anterior and posterior for each specific sample, and segmentation may then require significant manual input.

In addition to the possibility that more specific hippocampal volumetric measures may reveal stronger relationships with CRF and learning than the volume of the entire hippocampus, there is also evidence that other measures of hippocampal structure and function may serve as important mediators in the CRF-cognition relationship. For instance, Schwarb and colleagues (2017) found that hippocampal viscoelasticity mediated the relationship between CRF and relational memory performance in young adults. Viscoelasticity provides a measure of microstructural integrity of the hippocampus, which may be more sensitive to individual differences than volume measurements. Further, tissue density, measured via diffusivity, has also been found to be sensitive to changes in fitness in a sample of older adults (Kleemeyer et al., 2016). Importantly, changes in density were related to changes in hippocampal volume, which suggests both measures are sensitive to fitness changes. For our purposes of extending from other papers that have examined hippocampal volume and understanding the role of the hippocampus in the relationship between CRF and learning on a specific episodic task, volume measurements of the whole hippocampus were sufficient. Our findings do support future research that extends into sub-regions or systems of the medial temporal lobe as well as research that evaluates other measures of hippocampal microstructure, such as viscoelasticity and diffusivity.

Another limitation of this study is the cross-sectional nature. The strongest evidence for a relationship between CRF, hippocampal volume, and memory would come from a randomized controlled trial involving a structured exercise intervention known to increase CRF. Many such studies have shown evidence of a relationship between aerobic exercise and cognition (for review see A. F. Kramer & Colcombe, 2018; Voss et al., 2019), though none have utilized a task like the EAL task, which specifically targets aspects of hippocampal function that are critically important for accumulating episodic information over repeated occurrences. Nonetheless, our findings do align with previous evidence of exercise interventions resulting in increased brain structure, brain function, and cognition, as well as with other cross-sectional studies of CRF, hippocampal volume, and various aspects of memory (Erickson et al., 2009; Hayes et al., 2016). The current study provides an extension of previous findings by targeting memory processes that rely on hippocampal circuits that decline with age and are important for episodic memory. The cross-sectional design allowed us to establish a relationship between CRF and the EAL task. Based on this, the EAL task can be utilized in intervention studies to evaluate specific PA-and CRF-related changes in various parameters of learning.

Importantly, while CRF is determined in part by genetics, it is also influenced by frequency and intensity of PA. Thus, if CRF is modifiable, which can in turn influence the severity of age-related effects on the brain, individuals may be able to influence their personal trajectory by increasing PA, which would increase the possibility of preventing age-related decline. As PA remains difficult to objectively quantify, CRF serves as an important marker for physiological processes that may be induced by PA. Further, CRF has been shown to have a positive relationship with brain function independent of PA. Specifically, CRF is related to higher functional connectivity in networks that are diminished by age (Voss et al., 2016). Mechanistically, research supports that the health benefits of PA come from a variety of pathways, including increases in heart rate and repetitive muscle contraction and usage, all of which trigger a beneficial slew of neurotrophic pathways, decreased inflammation, improved body composition, and improved metabolic processes (for review see Warburton, Nicol, & Bredin, 2006). These pathways may also be systemically affected by CRF, such that individuals with higher CRF have elevated baseline activity of certain pathways and processes. It is also possible that differing levels of CRF, or other physiological characteristics, influence the extent to which PA acutely and chronically impacts the body and brain.

In sum, our findings support a relationship between CRF, hippocampal volume, and learning rate on a task that was specifically designed to engage hippocampal processes that decline with aging. We have shown that higher CRF predicts faster associative learning and that hippocampal volume serves as a biomarker of this relationship. The EAL task can be used in the future as a rich source of data that represents the acquisition speed of associative bindings, which provides insight about the integrity and function of hippocampal circuitry. The current study moves this field forward by combining strong measures of CRF and brain structure with a novel task that is sensitive to age and targets hippocampal-based learning.

## Supporting information

Acknowledgement

