## Supplementary materials for "Cardiorespiratory fitness predicts greater hippocampal volume and rate of episodic associative learning in older adults"

Supplemental materials

**Supplemental support for construct validity of the EAL task**

**Rationale**: A main goal of this study was to examine performance on learning tasks proposed to rely heavily on the hippocampus, which is known to experience age-related decline. In addition to the EAL task, we examined the Spatial Reconstruction task, which has previously been shown to require spatial relational memory (Watson, Voss, Warren, Tranel, & Cohen, 2013). The requirement of recalling the specific location previously associated with an object in relation to other objects in space should rely on the binding function of the hippocampus (Ryan, Lin, Ketcham, & Nadel, 2010). This type of ability is related to hippocampal volume in healthy adults (Monti et al., 2015) and is impaired in individuals with hippocampal amnesia (Konkel & Cohen, 2009; Watson et al., 2013). Specifically, Watson and colleagues (2013) found that hippocampal damage was associated with a disproportionate tendency to swap the location of two items. This finding demonstrated that the hippocampus plays a critical role in object-location binding in forming distinct inter-object relational memories. Another study of this task (Schwarb, Johnson, McGarry, & Cohen, 2016) found that in healthy young adults, the hippocampal characteristic of viscoelasticity (which reflects mictrostructural aspects of gray-matter structures, such as axonal connection integrity) was also positively correlated with better performance. Taken together, these studies show that the spatial reconstruction task has been well supported to critically rely on the structural integrity of the hippocampus, and can be used as a validated hippocampal-based task with which to evaluate and establish construct validity with our EAL task.

**Methods:**

*Spatial Reconstruction.* Twelve participants completed the iPad version of this task as described in Clark et al. (2018) which was originally developed by and described in Huttenlocher and Presson (1979). Briefly, participants were instructed to memorize the location of 5 novel “squiggle” objects on the screen. After 60 seconds of study and a 6 second delay, objects appeared at the bottom of the screen and participants used their finger to drag each object to its original location. Participants completed 30 trials.

The remaining 33 participants completed the computer version of the task, which was developed by Monti et al. (2015). The task was identical to the iPad version except that each trial included 6 objects instead of 5, and there were a total of 20 trials.

*Analysis.* The main outcome of interest for this task is the rate of swaps between objects. Rate was calculated as the number of swaps per trial divided by the total number of possible swaps (number of relations). For example, with 5 objects, there were 10 object pairs for possible swaps. With 6 objects, there were 15 possible swaps. Because the two versions of the task (iPad and computer) had different total numbers of objects, the distributions of swap rates were significantly different between the two samples (t = 4.3, p < .01). The average swap rate from the iPad sample (.06) was about half that of the computer version (.13) likely because the computer version consisted of 6 objects. For this reason, we normalized swap rate within each sample, and used the resulting z-score as a dependent variable.

**Results**

*Learning rate on EAL task correlates with performance on spatial reconstruction task.*

To determine construct validity of the EAL task, we tested whether learning rate on the EAL task correlates with a well-known relational memory metric, swap error rate on the SR task. EAL task learning rate was negatively correlated with swap error rate on the SR task (r = -0.45, p < .01), meaning individuals who learned at a faster rate on the EAL task also tended to make fewer swap errors than those individuals who learned more slowly on the EAL task. This suggests that a common relational process may underlie performance on both tasks. The rate of swap errors on the spatial reconstruction task was also negatively related to hippocampal volume (-.39, p < .01), such that individuals with larger hippocampal volume tended to make fewer swap errors.
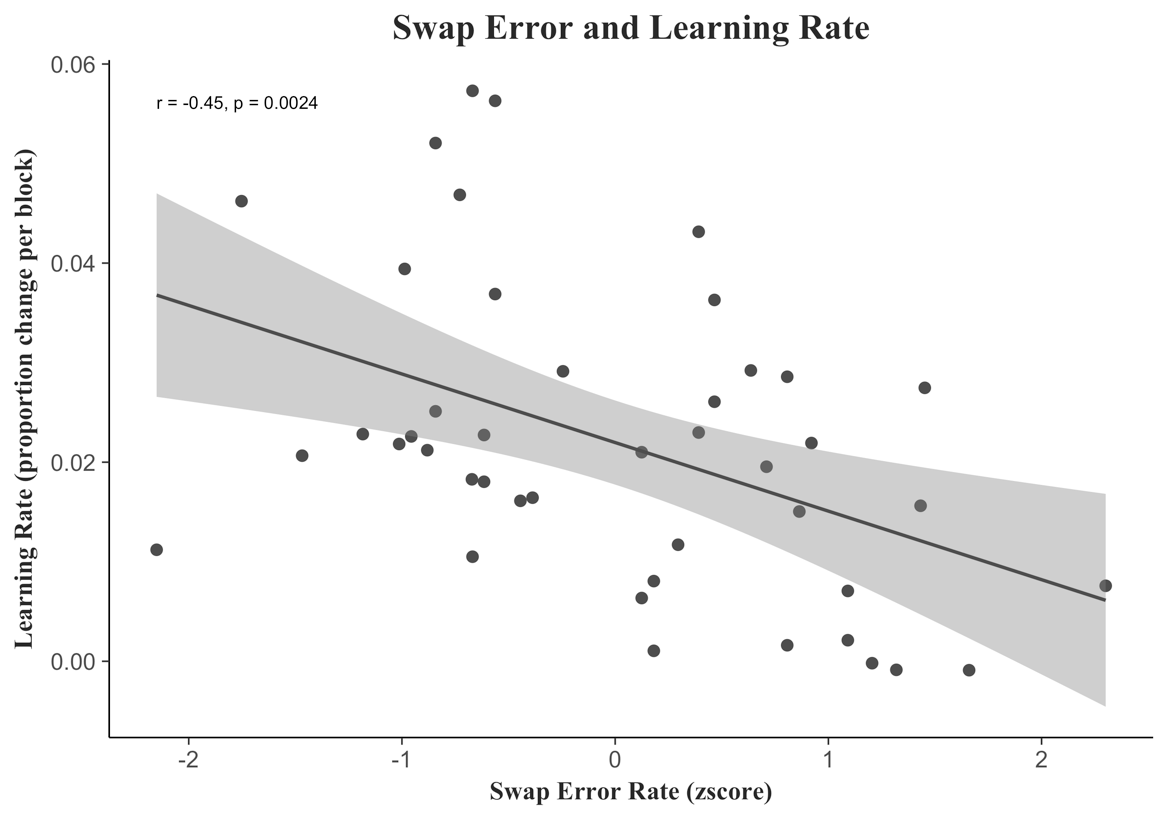

*CRF was not correlated with performance on the Spatial Reconstruction task*

CRF did not predict outcomes on the SR task (r = -.15, p = .3).

**Discussion**

We found support for construct validity of the EAL task, as performance was strongly related to swap error from the SR task, which has been shown to relate to multiple indices of hippocampus integrity and for which the hippocampus is critical as demonstrated by amnesiac patients (Monti et al., 2015; Schwarb et al., 2015; Watson et al., 2013). Interestingly, Schwarb and colleagues (2017) found that hippocampal tissue integrity, as measured by magnetic resonance elastography, was related to aerobic fitness and mediated the benefits of fitness on relational memory, as measured with a composite performance score from the SR task. In this study, hippocampal volume (a more gross measure of tissue structure than elastography) did not similarly mediate the relationship between fitness and memory. This relationship, however, was only tested and observed in young adults. It remains unknown whether hippocampal viscoelasticity would relate to aerobic fitness and relational memory in older adults. In our sample, hippocampal volume did correlate with SR performance, unlike the finding from Schwarb and colleagues (2017). As the authors noted, it may in fact be that hippocampal volume is adequately sensitive in older adults, whereas in younger adults, a more specific microstructural measure may be needed to evaluate the relationship between tissue integrity and relational memory. Nonetheless, future studies could evaluate elastography in older adults as well to determine if it is an even more sensitive measure than volume.

Clark, R., Hazeltine, E., Freedberg, M., & Voss, M. W. (2018). Age differences in episodic associative learning. *Psychology and Aging, 33*(1), 144-157. doi:10.1037/pag0000234

Huttenlocher, J., & Presson, C. C. (1979). The Coding and Transformation Information of Spatial. *394*, 375-394.

Konkel, A., & Cohen, N. J. (2009). Relational memory and the hippocampus: representations and methods. *Frontiers in neuroscience, 3*(2), 166-174. doi:10.3389/neuro.01.023.2009

Monti, J. M., Cooke, G. E., Watson, P. D., Voss, M. W., Kramer, A. R., & Cohen, N. J. (2015). Relating Hippocampus to Relational Memory Processing across Domains and Delays. *Journal of Cognitive Neuroscience, 27*(2), 1-10. doi:10.1162/jocn

Ryan, L., Lin, C., Ketcham, K., & Nadel, L. (2010). The role of medial temporal lobe in retrieving spatial and nonspatial relations from episodic and semantic memory. *Hippocampus, 20*(1), 11-18. doi:10.1002/hipo.20607

Schwarb, H., Johnson, C. L., Daugherty, A. M., & Hillman, C. H. (2017). NeuroImage Aerobic fi tness , hippocampal viscoelasticity , and relational memory performance. *NeuroImage, 153*(December 2016), 179-188. doi:10.1016/j.neuroimage.2017.03.061

Schwarb, H., Johnson, C. L., McGarry, M. D. J., & Cohen, N. J. (2016). Medial temporal lobe viscoelasticity and relational memory performance. *NeuroImage, 132*, 534-541. doi:10.1016/j.neuroimage.2016.02.059

Schwarb, H., Watson, P. D., Campbell, K., Shander, C. L., Monti, J. M., Cooke, G. E., . . . Cohen, N. J. (2015). Competition and cooperation among relational memory representations. *PLoS ONE, 10*(11), 1-22. doi:10.1371/journal.pone.0143832

Watson, P. D., Voss, J. L., Warren, D. E., Tranel, D., & Cohen, N. J. (2013). Spatial reconstruction by patients with hippocampal damage is dominated by relational memory errors. *Hippocampus, 23*(7), 570-580. doi:10.1002/hipo.22115
