## Supplementary material for "Cardiorespiratory fitness predicts greater hippocampal volume and rate of episodic associative learning in older adults": Acknowledgement

This research was supported by Grant 5R21AG048170 from the National Institutes of Health/National Institute on Aging. The authors thank Lauren Reist and the University of Iowa MRI Research Facilities (MRRF) technologists for their help with data collection.
